# The enhancement of activity rescues the establishment of *Mecp2* null neuronal phenotypes

**DOI:** 10.1101/2020.04.06.027995

**Authors:** Linda Scaramuzza, Giuseppina De Rocco, Genni Desiato, Clementina Cobolli Gigli, Martina Chiacchiaretta, Filippo Mirabella, Davide Pozzi, Marco De Simone, Paola Conforti, Massimiliano Pagani, Fabio Benfenati, Fabrizia Cesca, Francesco Bedogni, Nicoletta Landsberger

## Abstract

*Mecp2* deficiency, the gene responsible for Rett syndrome (RTT), affects brain maturation by impairing neuronal activity, transcription and morphology. These three elements are physiologically linked in a feed-forward cycle where neuronal activity modulates transcription and morphology to further increase network maturity. We hypothesized that the reduced activity displayed by maturing *Mecp2* null neurons during development could perturb such cycle, sustaining an improper transcriptional program that, ultimately, impairs neuronal maturation. Accordingly, we show that by enhancing activity within an early time window, Ampakine redirects, *in vitro*, the development of null neuronal networks towards more physiological routes. Similarly, the administration of the drug to newborn null offspring delays the progression of symptoms, significantly prolonging life span. Our data highlights the role of altered neuronal activity during the establishment of *Mecp2* null networks and the importance of such early defects to the typically poor maturity of RTT brain functions in adulthood. We propose the existence of an “early molecular phase” of Rett syndrome, a detailed description of which might disclose relevant targets for new rescue treatments.

## Introduction

Mutations in the X-linked Methyl-CpG-binding protein 2 (*MECP2*) gene are associated with a number of neurological conditions among which the most frequent is Rett syndrome (RTT; Amir *et al*, 1999). RTT is the most common cause of severe intellectual disability in females, as it affects one female every 10000 born alive (Chahrour & Zoghbi, 2007). Given the high levels of MeCP2 in the brain, the neurological features of RTT are by far the most thoroughly described. Smaller brain size and weight, thinner corpus callosum and reduced cortical thickness are typical of the pathology (Armstrong *et al*, 2001; Carter *et al*, 2008; Belichenko *et al*, 2009). Hemizygous *Mecp2* null male recapitulate much of these defects in adult stages and feature reduced life span (Guy *et al*, 2001). Defective neuronal features have been reported as well, as soma size, dendritic branching, number of spines and synaptic contacts are typically reduced (Guy *et al*, 2001; Bedogni *et al*, 2016; Baj *et al*, 2014; Belichenko *et al*, 2009; Chao *et al*, 2007; Fukuda *et al*, 2005; Kishi and Macklis, 2004; Sampathkumare *et al*, 2016; Rietveld *et al*, 2015).

Besides morphological alterations, impaired neuronal functions were also observed in adult mice, resulting in a complex derangement of brain activity (Nelson & Valakh, 2015). In particular, the absence of Mecp2 reduces the miniature excitatory postsynaptic currents (mEPSCs) but not the miniature inhibitory postsynaptic currents (mIPSCs), therefore leading to imbalance of the excitatory/inhibitory (E/I) ratio (Nelson *et al*, 2006; Tropea *et al*, 2009). This is in line with the reduced expression of the immediately early gene *cFos* in the forebrain, a surrogate of neuronal activity (Kron *et al*, 2012). Conversely, the lack of Mecp2 in the hippocampus causes a reduction in mIPSCs (Calfa *et al*, 2015; Katz *et al*, 2016).

The outbreak of symptoms typically follows an apparently normal developmental phase that lasts 6-18 months in humans (Cosentino *et al*, 2019) and roughly 40 days in *Mecp2* null animals (Guy *et al*, 2001; Chen *et al*, 2001). However, subtle symptoms in RTT girls have been described even before the onset of the overt phase of the pathology (Fehr *et al*, 2011; Dolce *et al*, 2013; Marschik *et al*, 2013; Cosentino *et al*, 2019). Similarly, animal models display several defects already during embryonic development (Bedogni *et al*, 2016; Cobolli Gigli *et al*, 2018; Mellios *et al*, 2018) or immediately after birth (Dani *et al*, 2005; Picker *et al*, 2006; Chao *et al*, 2007). Such evidence demonstrates that the lack of MeCP2 affects early development of the mammalian brain although the phenotypic consequences of it become overt later on (Ip *et al*, 2018; Cosentino *et al*, 2019).

In line with the role of Mecp2 as master regulator of transcription (Skene *et al*, 2010; Lagger *et al*, 2017; Shah & Bird, 2017), the emergence of transcriptional maturity is affected in null samples both *in vivo* (Cobolli Gigli *et al*, 2018) and *in vitro* (Mellios *et al*, 2018) and the ability of null neurons to respond to external *stimuli* is reduced (Bedogni *et al*, 2016). We thus hypothesized that the poor functionality of null neurons during development could be a key element to explain the thoroughly described poor maturity of adult *Mecp2* deficient neuronal networks. Indeed, while neuronal activity is surely involved in the maintenance of mature synaptic structures (Greer & Greenberg, 2008; Flavell & Greenberg, 2008), early patterns of spontaneous activity have been reported already during embryonic and early postnatal cortical development (Corlew *et al*, 2004; Platel *et al*, 2005; Spitzer, 2006). However, such spontaneous activity cannot be simply considered a measure of the gradually increasing maturation of membrane excitability, as it also drives neurodevelopmental processes that include refinement of neuronal identity, newborn neurons migration, survival and connectivity (Luhmann *et al*, 2015; Yamamoto and Lopez-Bendito, 2012; Bonetti and Surace, 2010; Weissman *et al*, 2004). In fact, neuronal activity is required at all developmental stages; in particular, electrical activity is thought to refine signaling via gene expression, providing checkpoints to validate or modulate genetic programs essential for the establishment of proper neuronal maturity (Spitzer, 2006). We thus tested the possible causative link between poor maturity and reduced neuronal activity in null samples by pharmacologically stimulating null neurons within early time windows. To focus on the earliest steps of neuronal maturation, we used neurons differentiated from cortical neuroprecursors (NPCs), which recapitulate the *in vivo* process of neuronal differentiation and the subsequent steps of gliogenesis (Paridaen & Huttner, 2014). Functional *in vitro* assessments were produced using differentiated neurons or primary neuronal cultures. Eventually, we validated our results *in vivo* using *Mecp2* null animals.

Our *in vitro* data demonstrate that by enhancing activity, Ampakine CX546 rescues *Mecp2* null neurons morphological and transcriptional defects and restores impaired neuronal network functions. Accordingly, our *in vivo* data show that the exposure of *Mecp2* null animals to CX546 within an early and short postnatal time window prolonged life span and ameliorated their behavioral scoring. Importantly, both *in vivo* and *in vitro* rescue effects lasted long after the exposure to the drug, implying the long persistence of such effects. Our results suggest that the enhancement of neuronal activity in *Mecp2* null neurons sets in place plasticity mechanisms that mitigate maturation defects likely through the modulation of the mentioned feed-forward cycle (Spitzer, 2006). We suggest that the time frame in which such mechanisms occur highlights an important early molecular phase of the pathology, during which null neurons are affected by a number of transcriptional derangements that we believe are prodromal of the subsequent onset of overt symptoms. Therapeutic strategies acting within this early molecular phase might be more effective in readdressing the pathological trajectory of null neurons development towards more physiological directions than later treatments.

## Results

### Lack of Mecp2 does not affect the early commitment of *in vitro* cultured cortical NPCs

To focus on early mechanisms of maturation of *Mecp2* null neuronal networks, we collected neuroprecursor cells (NPCs) from E15 cerebral cortex and induced their *in vitro* differentiation as previously described (Magri *et al*, 2011; Cobolli Gigli *et al*, 2018). Being synchronously generated, the stage of differentiation of such populations is roughly comparable. Defects masked in heterogeneous populations are consequently easier to detect, an advantage in the case of Mecp2 studies, where in most cases defects are very subtle (Bedogni *et al*, 2014). Moreover, this model enables to reduce the use of mice and is suitable for pharmacological manipulations (Gorba & Conti, 2013).

NPCs spontaneously proliferate in a self-renewing manner in the presence of the mitogens EGF and FGF2 (Gritti *et al*, 1999), while the presence of fetal bovine serum drives their differentiation into three main cellular populations of the brain: neurons, astrocytes and oligodendrocytes (Figure 1A; Magri *et al*, 2011). We measured the distinct cellular populations at different developmental stages of NPCs differentiation by immunostaining wild type (wt) and *Mecp2* null cultures for Tuj1, Gfap and Olig2. We found that the lack of Mecp2 did not affect the percentage of neurons, astrocytes and oligodendrocytes, respectively (Figure 1B-J) at any differentiation stage, in line with previous observations (Kishi & Macklis, 2004). In particular, neurons contributed for roughly 27,2±1,1 and 22,2±0,6 at DIV8 and for 21,6±1,8 and 19,7±1,6 at DIV22 in wt and null samples respectively (Figure 1D). The number of Gfap positive cells increased over time, spanning from roughly 35% at DIV8 to 70% at DIV22 in both genotypes, therefore explaining the reduction in the percentage of neurons from DIV8 to DIV22 (Figure 1G). Gfap positive cells were, therefore, generated at a later time point than neurons, in line with the dynamics of glial cells genesis and differentiation *in vivo* (Paridaen & Huttner, 2014). Similarly, the percentage of oligodendrocytes (roughly 7%) remained equal throughout differentiation (Figure 1J).

**Figure 1:**
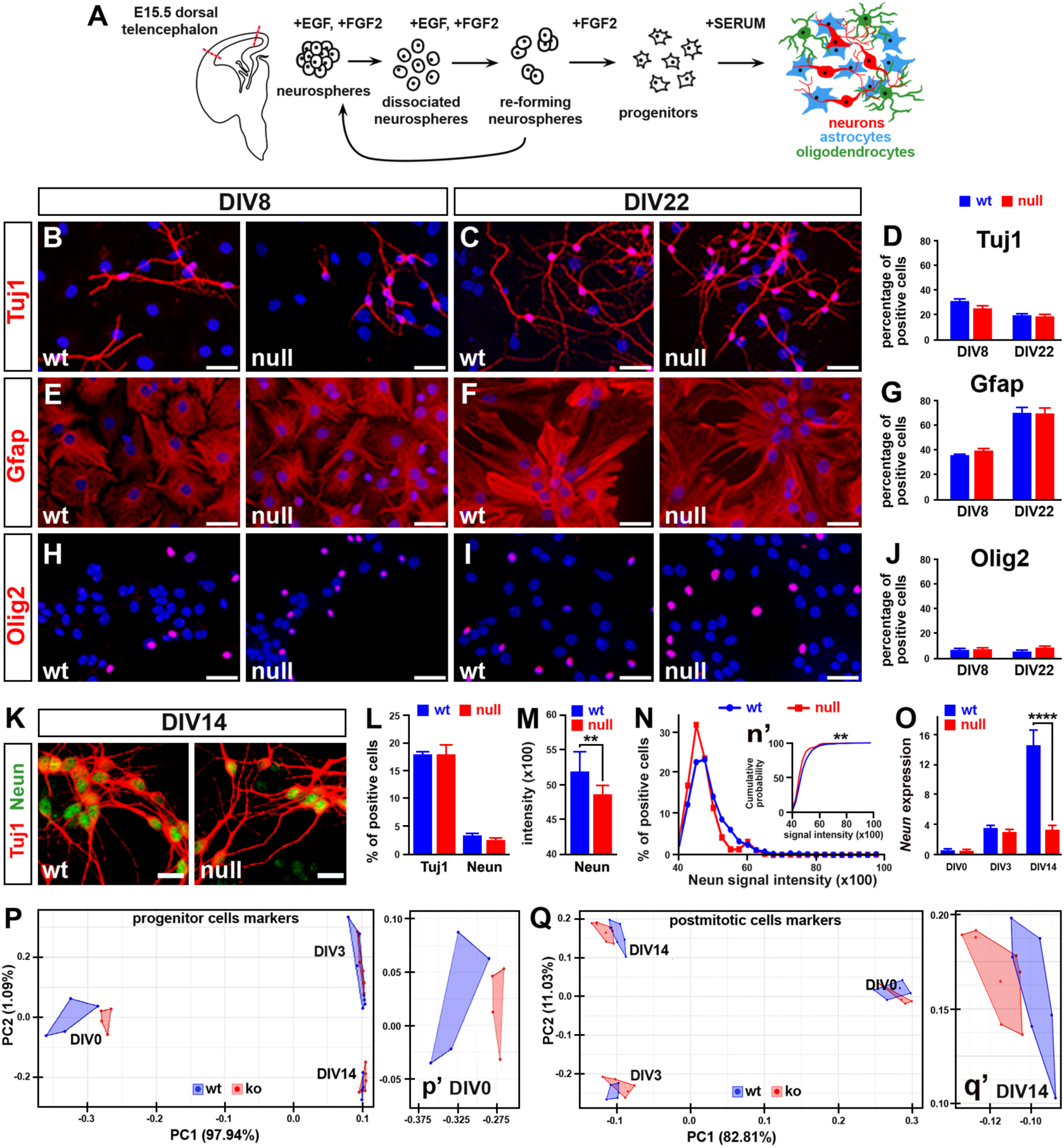
Lack of Mecp2 does not affect the commitment of NPCs but delays their differentiation. **A**: Schematic representation of the differentiation protocol and the cell types composing cultures differentiated from NPCs (adapted from Carlessi *et al*, 2013). **B-J**: Representative immunofluorescences of the cell types composing the WT and *Mecp2* null cultures differentiated from NPCs. Antibodies against Tuj1, Gfap and Olig2 were used to recognize neurons, astrocytes and oligodendrocytes. The percentage of cells positive for each marker was counted over the total of DAPI positive cells (100%). Histograms in D, G and J show the mean percentage ± SEM of positive cells. Sample size is n ≥3 wells obtained from two independent preparations. No difference between genotypes was highlighted (Two-way ANOVA); the variation through time resulted significant for the levels of Tuj1 and Gfap (Two-way ANOVA: F(1,16)=9.542, p-value<0.05 and F(1,10)=68.95, p-value<0.0001 respectively). **K**: Representative immunostaining of Neun (green) and Tuj1 (red) positive neurons at DIV14. **L and M**: Histograms show the mean percentage ± SEM of Neun and Tuj1 positive cells and the average Neun intensity in wt and null samples. The percentage of cells positive for each marker was counted over the total of DAPI positive cells. Student’s t-test highlighted a significant decrease of Neun intensity in null samples. **: p-value<0.01. Sample size is n ≥5 wells obtained from two independent preparations. **N and n’**: The binned distribution of Neun intensity was plotted as both frequency distribution graphs, in panel N, and cumulative probability plot in panel n’. The null neurons distribution is significantly shifted towards low levels of Neun intensity, Kolmogorov-Smirnoff test**: p-value<0.01. Each value in panel N is represented as mean ± SEM obtained from two different preparations measuring 900 cells. **O**: *Neun* expression levels assessed at DIV0, DIV3 and DIV14 are represented as 2^-Δct^. Histograms show the mean ± SEM from at least 5 samples obtained from two independent preparations. Two-way ANOVA indicated both a significant genotype effect (F(2,28)=19,60; p<0,0001) and time effect (F(2,28)=93,70; p<0,0001). Bonferroni post-hoc test revealed significant differences between genotype at DIV 14 but not at DIV0 and DIV 3. ****: p-value<0.001. **P-Q**: PCA analyses show the distance between the transcriptional identities of wt and null samples using different sets of genes (see Table S1). 16 genes typically expressed by cortical neuroprogenitors were considered in P; in Q, 51 genes typically expressed by neurons. P’ and q’ are close ups at different DIVs. Three independent preparations were used to produce the points composing the wt and null cluster. Each point represents a single well. Scale bars: B-J: 30 µm; K: 15 µm.

At DIV8 we detected cells there were not positive for any of the three markers but were positive for Nestin, which suggested the persistent presence of proliferating cells at that stage (Figure S1A). Such population eventually disappeared at DIV22 (Figure S1B), thus following the decrease in the expression of the cell cycle marker *Ki67* over time (Figure S1C). We conclude that at DIV22 most cells terminally differentiated into their specific fate with no influence of the genotype on the final commitment.

To further support these findings, we repeated the analysis on NPCs derived from heterozygous *Mecp2*^*-/+*^ female cerebral cortices, which are generally subjected to random X chromosome inactivation (Figure S2A-D). Consistent with our previous results (Figure 1D and 1G), at DIV22 we found no different percentage of either neurons (Tuj1^+^) or glial cells (Gfap^+^) between the Mecp2 positive and Mecp2 negative populations (Figure S2B-D).

### Lack of Mecp2 delays the transcriptional maturation of *in vitro* differentiated NPCs

To verify whether our model recapitulated the previously described delayed mechanisms of neuronal maturation (Mellios *et al*, 2018; Cobolli Gigli *et al*, 2018; Bedogni *et al*, 2016), we exploited the expression levels of Tuj1 and Neun, a neuronal-specific marker that increases with neuronal maturation at a slower rate compared to Tuj1 (Walker *et al*, 2007; Menezes *et al*, 1994). We analyzed null and wt cultures at DIV14 (Figure 1K-O) and while the number of Neun and Tuj1 positive cells overlapped between wt and null samples (3,2± 0,4 and 2,5± 0,3; 1L), the intensity of the Neun immunosignal was significantly decreased in null samples (Figure 1M), as well as the percentage of cells expressing low levels of it (Figure 1N and n’).

Moreover, in wt cultures the mRNA levels of *Neun* increased with time, thus following neuronal maturation, while, in the case of *Mecp2* null cultures, the expression of *Neun* did not change between DIV3 and DIV14, thus highlighting a delay or arrest of transcriptional maturation (Figure 1O). Prompted by this evidence, we next measured the transcription of two sets of genes that included respectively markers of neuronal progenitors and post-mitotic neurons (Table S1). To visualize the segregation of wt and *Mecp2* null samples we used Principal Component analysis (PCA). The use of the set of 16 genes typically expressed by NPCs (Figure 1P) unmasked a clear segregation between the two genotypes at DIV0 (Figure 1p’) that disappeared at later time points. This is in line with the fact that neurogenesis is delayed, but not compromised in null samples (Cobolli Gigli *et al*, 2018). Conversely, the use of a set of 51 genes expressed by post-mitotic neurons revealed an opposite situation in which the overlapping transcriptional identities at DIV0, segregated at DIV14 (Figure 1 Q-q’). Interestingly, when we analyzed the significance of such transcriptional changes through two-way ANOVA, we noticed an overall downregulation of the selected genes and, in particular, we found decreased levels of several components of the neuronal activity machinery, including receptors, ionic channels and intracellular molecules (Figure S3). Altogether, these results confirmed that our *in vitro Mecp2* null model recapitulates the maturation delay already described *in vivo* during corticogenesis (Cobolli Gigli *et al*, 2018). Moreover, the deregulation of genes known to be important for neuronal responsiveness to external stimuli not only confirmed previous *in vivo* results, but also suggested an impaired neuronal activity of *Mecp2* null NPC derived neurons, as already reported in null primary neurons (Bedogni et al.2016).

### The *in vitro* established *Mecp2* null neuronal networks are functionally and morphologically immature

To verify the impact of maturation defects on the activity of *Mecp2* null neuronal networks, we next focused on functional assessments. We plated dissociated NPCs at high density (8000 cells/µl) on Multi Electrode Arrays (MEAs) chips to record spontaneous electrical activity (Figure 2A and B). The same population of cells was recorded at two time points (DIV18 and 22), to monitor the functional development of cell-to-cell communication and the generation of an active network over time.

**Figure 2:**
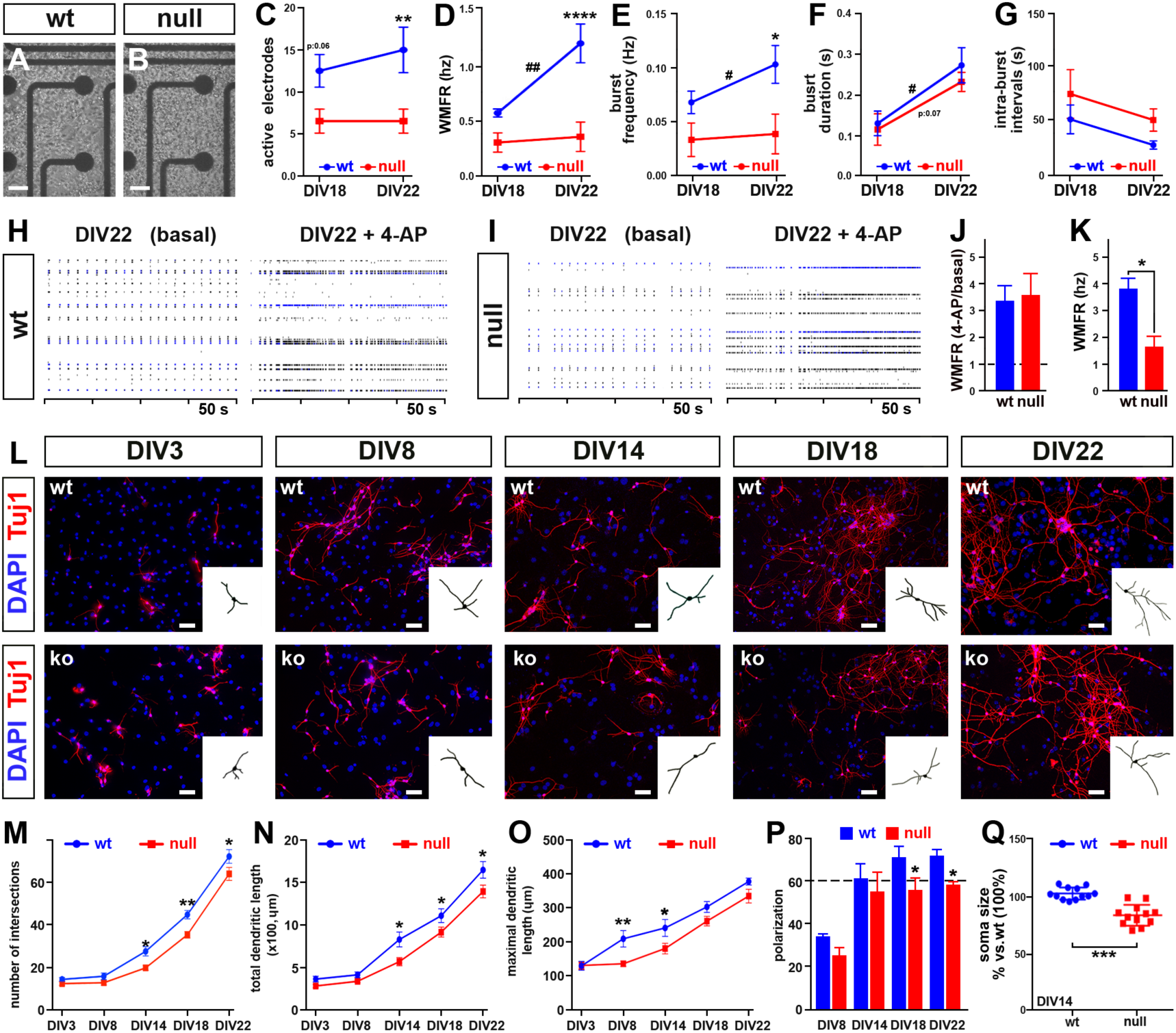
Neurons derived from *Mecp2* null NPCs are morphologically more immature than wt controls and establish less active networks. **A**,**B**: Wt or *Mecp2* null NPCs plated on the grid of 64 MEA electrodes. **C-G**: Line graphs represent the changes through time of: number of active electrodes, Weighted Mean Firing Rate (WMFR), burst frequency, burst duration, intra-burst intervals. Difference between genotype and through time were analyzed using Two-way ANOVA for repeated measures and Bonferroni post-hoc test; * refers to differences between genotypes (*: p-value<0.05; **: p-value<0.01; ***: p-value<0.005); # refers to differences between the two time points (#: p-value<0.05; ##: p-value<0.01). **H-K**: Raster plots (H and I) and their quantitation under basal conditions and upon 100µM 4-AP exposure. The magnitude of the response after stimulation was expressed either as fold change (J) or as the averaged maximal values reached after 4-AP exposure (K). Dotted line in J represents untreated controls (set to 1). Values are represented as mean ± SEM. Student’s t-test, *: p-value<0.05. Sample size (Panel A-K): n=8 wells deriving from three independent preparations. **L**: Wt and null neurons counterstained with an antibody against Tuj1 and DAPI at different time points. The binary mask is representative of the neurons used to study morphology. **M-Q**: Each graph represents a different morphological parameter analyzed at selected time point. Two way ANOVA indicated both a significant time effect ((F 4,20)=282,5 and p<0,0001 for the number of intersection; (F 4,26)=104,5 and p<0,0001 for the total dendritic length; (F 4,20)=58,51 and p<0,0001 for the maximal dendritic length; (F 3,22)=38,32 and p<0,0001 for polarization) and genotype effect ((F 1,20)=25,1 and p<0,0001 for the number of intersections; (F 4,26)=16,0 and p=0,005 for the total dendritic length; (F 1,20)=17,24 and p=0,005 for the maximal dendritic length; (F 1,22)=18,76 and p<0,003 for polarization). The dotted line in panel P represents the cutoff above which a culture is defined polarized (Horton *et al*, 2006). Values are represented as mean ± SEM. *: p-value<0.05; **: p-value<0.01; ***: p-value<0.005. Sample size (Panel M-Q): n=3 wells deriving from three independent preparations. Scale bar: 50 µm (A and B); 30 µm (L).

The spontaneous activity of the developing networks was assessed through five parameters, i.e. number of active electrodes, weighted mean firing rate (WMFR), frequency and duration of spontaneous bursts, and intra burst intervals (Figure 2C-G). The maturation of wt networks was demonstrated by the significant increase of WMFR, burst frequency and burst duration at DIV22 compared to DIV18 and the increasing trend in the number of active electrodes in the network, in line with published observations on the same model (Groot *et al*, 2014). The lack of Mecp2 instead drove a markedly alteration of this pattern, with null cultures showing a lack of functional maturation between DIV18 and DIV22, as indicated by the analysis of active electrodes, WMFR, burst frequency and duration (Figure 2C-F). The intra-burst intervals showed a similar trend between the two genotypes (Figure 2G). These data suggest that, differently from wt, null neurons do not integrate into an increasingly cohesive network, indicative of a defect in neuronal maturation. Noticeably, the parameters describing network activity in our model were lower in magnitude compared to primary neurons, as expected given the different level of maturity displayed by the two *in vitro* models (Chiacchiaretta *et al*, 2017).

After measuring spontaneous activity, we studied the evoked response upon exposure to 100 µM 4-AminoPiridine (4AP) at DIV22 (Figure 2H-K). Although the fold increase in WMFR was comparable between the two genotypes (Figure 2J), when considering the absolute WMFR values, the response was 2-fold higher in wt samples compared to null samples (2,9±0,4 in wt and 1.5±0,6 ko samples; Figure 2K). These results demonstrate that *Mecp2* null networks established from differentiated NPCs are basally less active compared to wt, in line with previous data (Bedogni *et al*, 2016; Kron *et al*, 2012; Katz *et al*, 2016; Sceniak *et al*, 2016).

We next analyzed the growth of dendritic branches along NPC development. The dendritic complexity analyzed in wt and *Mecp2* null neurons increased through time in both genotypes (Figure 2L). However, starting from DIV 14, null neurons exhibited a poorer dendritic growth compared to wt, as we detected a significant reduction of both intersections number and total dendritic length (Figure 2M,N), while maximal dendritic length appeared reduced at earlier time points (Figure 2O). Similarly, the percentage of polarized null neurons resulted significantly diminished from DIV18 (71% and 55,8% in wt and null neurons respectively, Figure 2P; see materials and methods for definition of polarized neurons). Eventually, in line with previous findings (Bittolo *et al*, 2016; Sampathkumar *et al*), NPC derived null neurons at DIV14 showed a reduction in soma size 2016; Figure 2Q).

Overall, these functional and morphological data suggest that our model reproduces many of the typical maturation defects generated *in vivo* and *in vitro* by the absence of Mecp2 (Bedogni *et al*, 2016; Baj *et al*, 2014; Rietveld *et al*, 2015; Sampathkumar *et al*, 2016).

### The enhancement of glutamatergic transmission within an early time window produces rescue effects that persist at later time points

Our hypothesis is that reduced activity-dependent signaling in *Mecp2* null neurons contributes to their defective transcriptional and morphological immaturity, further reinforcing a defective response to stimuli. We thus studied whether a pharmacological enhancement of neuronal excitability could rescue the observed maturation defects. With this aim maturing neurons were treated with Ampakine CX546, a positive modulator of glutamatergic transmission that slows down the deactivation rate of AMPA receptors without causing desensitization (Lynch *et al*, 2006). Initially, we compared the expression of the AMPA receptor subunit 2 (*Gria2*), one of the main targets of CX546, in the two genotypes at different developmental stages. We found that wt and *Mecp2* null neurons expressed similar amounts of the subunit at DIV3 and 7, while at DIV14 null neurons showed a significantly reduced expression compared to wt (Figure S4A). Based on published data, we tested at first two different doses of CX546 (10 and 20 µM; Schitine *et al*, 2012). In both cases, daily addition from DIV7 to DIV10 did not produce toxic effects (Figure S4B), while chronic exposure to 20 µM CX546 (from DIV1 to DIV10) resulted in only 50% cell survival (Figure S4C). We thus designed a protocol in which 20 µM CX546 was added every day to the cultures for four days and then washed out. To assess whether different time windows of CX546 exposure could produce different rescue effects, CX546 was added either from DIV3 to DIV6 or from DIV7 to DIV10 (Figure 3A) followed by washout. Samples were then collected for morphological and transcriptional assessments at DIV14. As shown in Figure 3B-C, CX546 treatments produced a significant amelioration of several morphological parameters in null neurons, including dendritic length, number of neurite intersection and soma size. Notably, the treatment resulted slightly stronger when the drug was early administered.

**Figure 3:**
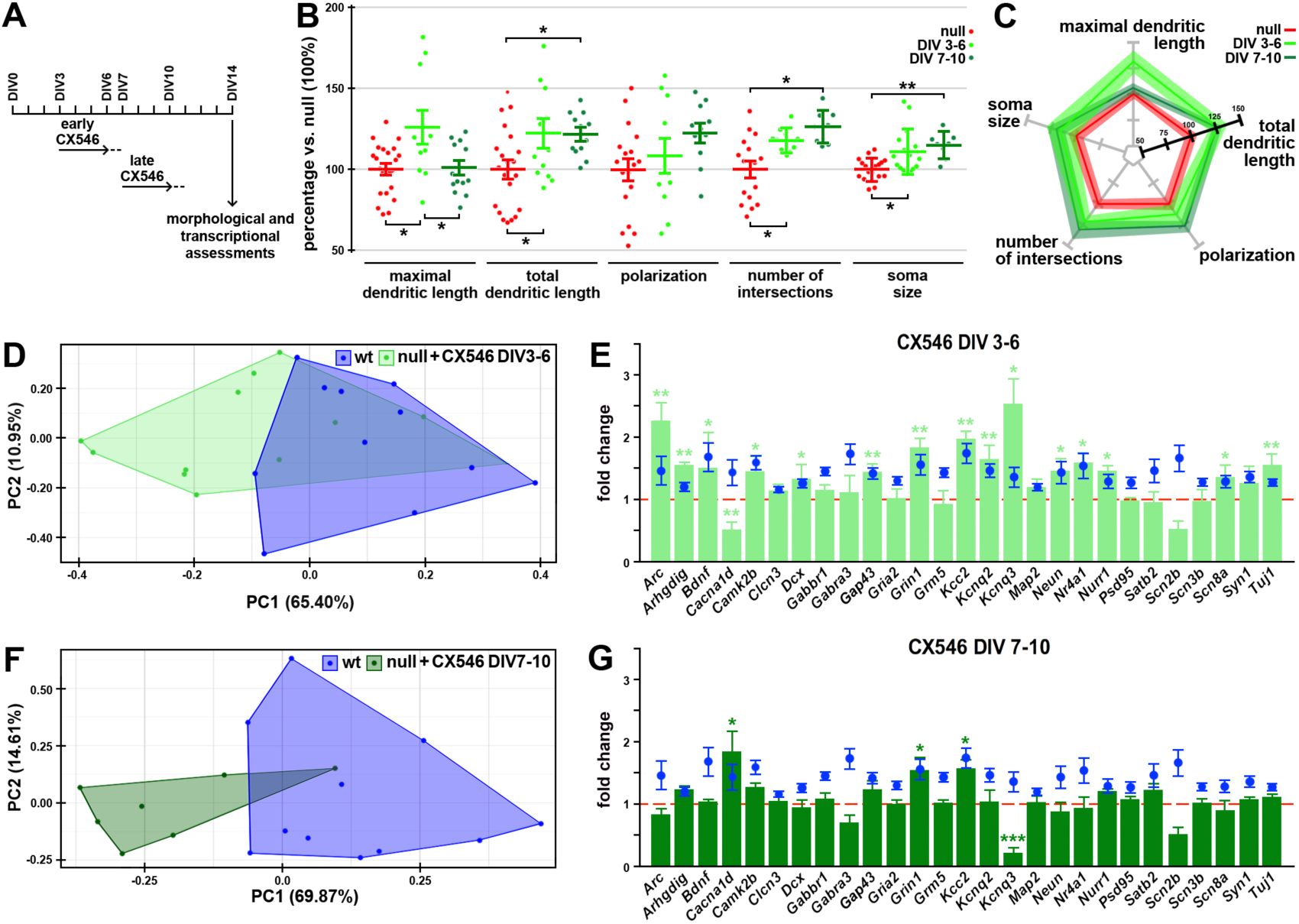
Early exposure to CX546 rescues morphological and transcriptional defects of null neurons. **A**: Schematic representation of the CX546 exposure. The drug was added to the cell medium from DIV3 to DIV6 and from DIV7 to DIV10; a complete wash-out of the medium was performed at the end of each treatment (DIV7 and DIV11 respectively). **B**,**C**: Five morphological parameters were measured in untreated and treated null and wt neurons at DIV14. One-way ANOVA indicated a significant treatment effect for the maximal dendritic length (F=6,306;p=0,0039), total dendritic length (F=4,880;p=0,0126), number of intersections (F=10,70; p=0,0003) and soma area (F=7,721;p=0,0015). The Tukey’s multiple comparison test highlighted a significant rescue for both the two treatments compared to the untreated group for the following parameters: number of intersections, total dendritic length, soma area. However, for the maximal dendritic length only the early treatment was statistically significant compared to untreated ko. *: p-value<0.05; **: p-value<0.01; ***: p-value<0.005. The variation of each of the selected parameters was summarized in the radar plot (C). Sample size (panels B,C): n≥18 wells for ctrl group and n≥10 wells for the treatment groups deriving from three independent preparations. Each point is representative of a single well. **D-G**: The two PCA analyses were based on the 43 genes selected in Figure S3 and highlighted the transcriptional differences between wt (in blue) and ko CX546 treated (light and dark green for early and late treatment respectively) samples. Bar graphs represent the expression values of the 27 selected genes after CX546 administration in the two different time windows (early in E and late in G). One way ANOVA followed by Tukey’s multiple comparison test was used to compare the expression of each gene between *Mecp2* null early and late treated samples and their corresponding untreated controls (set to 1, red dotted line). Wt levels are depicted in blue. Values are represented as mean ± SEM. *: p-value<0.05; **: p-value<0.01. Sample size (panels D-G): n≥8 wells deriving from two independent preparations.

Next, we assessed transcription on RNA samples prepared from wt controls and null neurons treated with either Ampakine or vehicle and collected at DIV14 (Figure 3D-G). Through PCA, we focused only on those transcripts that resulted unaffected by any batch effect (43 genes; Figure S3; see materials and methods). When we analyzed the transcriptional profile after early CX546 exposure (DIV3-DIV6), we found that the distance from wt and null treated identities was reduced compared to the basal clusters segregation observed in Figure 1Q and q’ (Figure 3D). In line with this, a detailed representation of the 27 transcripts selected for their statistically significant down-regulation between wt and null samples, showed that 15 of them were increased to levels that were significantly higher compared to untreated null samples (red dotted line in Figure 3E) and comparable to those of wt control (in blue in Figure 3E).

Importantly, CX546 was more effective when added from DIV3 to DIV6 compared to DIV 7-DIV10, as shown by PCA analysis (Figure 3D-F), where the late treatment was not able to produce a significant transcriptional overlap between null treated and wt control samples. Accordingly, among the selected 27 genes only 3 were significantly increased (Figure 3G).

We next tested the effectiveness of CX546 on primary neuronal cultures produced from null and wt E15 embryonic cortices, a model that has been thoroughly used to highlight functional defects driven by the lack of Mecp2. In line with our previous observations (Bedogni *et al*, 2016), primary null neurons display a lower neuronal responsiveness to extracellular stimuli, as indicated by the analysis of intracellular Ca^2+^ transients induced by 100 µM NMDA, with *Mecp2* null primary cultured neurons showing a significantly reduced responsiveness compared to wt at DIV14 (Figure 4B-C). The two bell shaped curves representing the distribution of the responses of wt and null samples in fact do not overlap, with the null curve shifted towards lower responses. However, the early exposure of null neurons to CX546 led the representing curve of null neurons (Figure 4B) to overlap with that of wt, therefore proving that the treatment was able to rescue the reduced responsiveness to NMDA. On the contrary, the late treatment with CX546 did not modify the null curve (Figure 4C), therefore indicating the higher effectiveness of the early treatment.

**Figure 4:**
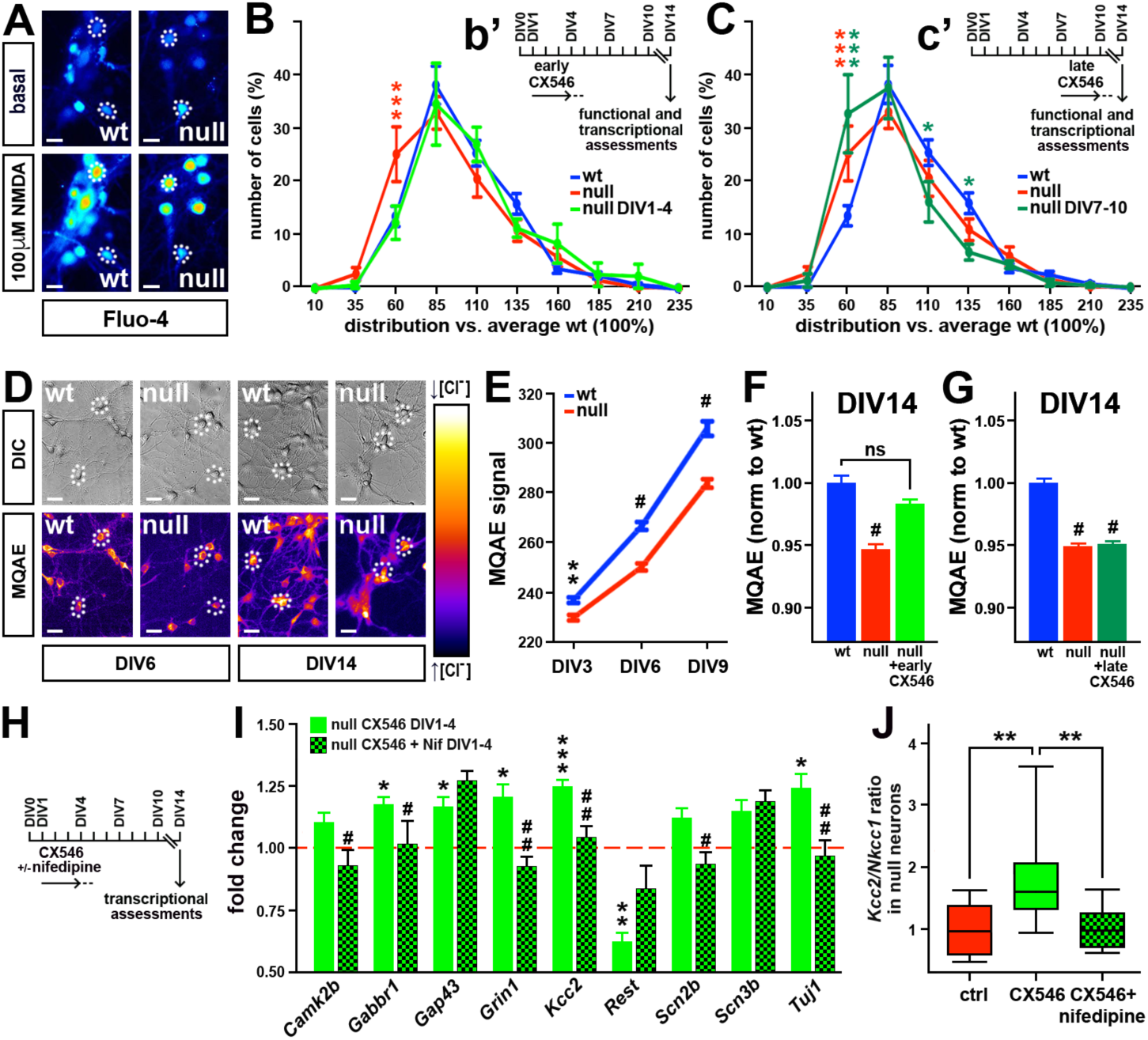
*Mecp2* null neuronal network responsiveness is rescued by early treatment with CX546. **A:** Representative images of DIV14 wt and *Mecp2* null primary cortical neurons loaded with Fluo-4 before and after NMDA 100 µM exposure. High fluorescent signals correspond to high calcium levels. Dotted cycles are representative of the Region Of Interest (ROI) used during the analysis. **B-C:** The graphs represent the binned distribution of the magnitude of NMDA induced calcium transients in wt (in blue), *Mecp2* null (in red) and treated *Mecp2* null (light and dark green for early and late treatment respectively). The distribution of responses on X-axis are expressed as percentage compared to wt (100%), values on Y-axis refer to percentage of cells over total (100%). Two way ANOVA followed by Bonferroni post hoc multiple comparison test was used to compare calcium responses between wt, *Mecp2* null and *Mecp2* null neurons early (B) and late treated (C) samples. *: P-value<0.05 ***: p-value<0.005. Red *: comparison between wt and ko; green *: comparison between wt and treated ko. B’ and c’ inserts represent the protocol of CX546 exposure; the dotted line indicates the wash out of the drug. Values are represented as mean ± SEM. Sample size (panels B,C): n= 11 and 10 animals for wt and *Mecp2* null control groups and n= 4 and 6 animals for early and late Ampakine treated groups. **D:** Representative images of DIV 6 and DIV14 wt and *Mecp2* null primary cortical neurons loaded with MQAE dye. High levels of fluorescence correspond to low level of intracellular chloride. Dotted cycles are representative of the Region Of Interest (ROI) used during the analysis. **E:** Line graphs represent intracellular chloride concentration at different consecutive time points in wt and *Mecp2* null neurons. Two way ANOVA indicated a significant effect for both time (F(2,1541)=840,9 and p<0,0001) and genotype (F(1,1541)=155,7 and p<0,0001). Bonferroni post hoc test revealed a statistically significant reduction of *Mecp2* null MQAE intensity at all the time points analyzed. **: p-value<0.01; #:p<0,0001. Values of MQAE intensity are represented as mean ± SEM. Sample size: n≥200 cells from three different embryos *per* genotype. **F,G**: Bar graphs representing the average values of MQAE intensity for early (F) or late (G) treated null neurons and their relative controls. One-way ANOVA followed by Tukey’s multiple comparison test was used to compare null control neurons and null treated neurons to wt neurons. #:p<0,0001. Values of MQAE intensity are represented as mean ± SEM. Sample size: n≥300 cells from five different embryos *per* genotype. **H:** Schematic representation of CX546 exposure, dotted line indicates drug wash out. **I-J**: In I One way ANOVA was used to compare the expression of 9 genes, selected among the ones analyzed in Figure 1, between *Mecp2* null primary neurons (set to 1, red dotted line), null neurons treated with CX546 from DIV1 to DIV4 and null neurons treated with CX546 and Nifedipine from DIV1 to DIV4. * refers to differences between *Mecp2* null untreated neurons and *Mecp2* null neurons treated with CX546 (*: p-value<0.05; **: p-value<0.01; ***: p-value<0.005); # refers to differences between *Mecp2* null neurons treated with CX546 and null neurons treated with both CX546 and Nifedipine (#: p-value<0.05; ##: p-value<0.01).In panel J we calculated the ratio between the expression level of *Kcc2* and *Nkcc1* (expressed as 2^-Δct^). One way ANOVA was used to compare the effects of the different treatment on null neurons compared to the untreated ones. Values are represented as mean ± SEM. *: p-value<0.05; **: p-value<0.01; ***: p-value<0.005. Sample size: n≥3 wells from four different embryos. Scale bars in A: 20 µm, in B: 20 µm.

Next, we asked whether other defective functional features of null neurons could be rescued by this treatment. To this aim we focused on the excitatory-to-inhibitory developmental GABA switch. A crucial parameter that participates to this process is the intracellular level of chloride, which progressively diminishes throughout neuronal development, enabling the transition of the GABA responses from excitation to inhibition (Ben-Ari *et al*, 2012). The molecular mechanisms driving these events are affected by the lack of Mecp2, resulting in a higher intracellular chloride level in mature null neurons (Tang *et al*, 2016). In line with this finding, the intracellular chloride, imaged at single cell level with MQAE dye, was higher in null neurons compared to control at each time point we analyzed (Figure 4D and E; the MQAE signal is inversely proportional to the chloride content).

Once again, the early exposure of null neurons to CX546 restored an intracellular chloride level comparable to wt neurons (Figure 4F), while no effect was observed in the late time window of treatment (Figure 4G). By testing the ability of CX546 to rescue the expression of genes typically affected by the absence of Mecp2, we confirmed the positive effect of the early treatment on transcription. Furthermore, by measuring the expression of the same genes in cells treated simultaneously with CX546 and Nifedipine, a blocker of L-type Calcium channels, we further validated our hypothesis by which the enhancement of activity drives the phenotypic maturation in *Mecp2* null neurons. Indeed, the early Ampakine treatment (from DIV1 to DIV4; Figure 4H-I) induced the expression of several genes (*Gabbr1, Gap43, Grin1, Kcc2* and *Tuj1*) that are otherwise downregulated in ko neurons (Fig.S3); this significant increment was generally not observed in the presence of Nifedipine, therefore confirming the involvement of voltage gated L-type Calcium channels in the mechanism of CX546 action. Similarly, Nifedipine abolished the reduction driven by CX546 on the expression of Rest, a gene that is generally upregulated in null samples (Abuhatzira et al, 2007). In good accordance with the functional data, in panel J we found that only CX546 increased the ratio of the expression levels of *Kcc2* and *Nkcc1*, therefore indicating a faster rate of maturation of GABA signaling (Tang *et al*, 2019; Hinz *et al*, 2019).

### Early treatment with Ampakine ameliorates behavioral deficits of *Mecp2* null mice

Prompted by the rescue effects driven by the early *in vitro* exposure to CX546, we assessed its effectiveness *in viv*o. To this aim, we used the CD1 *Mecp2*^tm1.1Bird^ RTT mouse model, which we recently characterized (Cobolli Gigli *et al*, 2016). *Mecp2* null and wt mice were injected subcutaneously daily with either CX546 (40mg/kg) or vehicle from P3 to P9 and the validity of the treatment was evaluated starting from P30 testing multiple readouts (Figure 5A). A total of 25 wt animals and 25 KO animals were used for this experiment.

**Figure 5:**
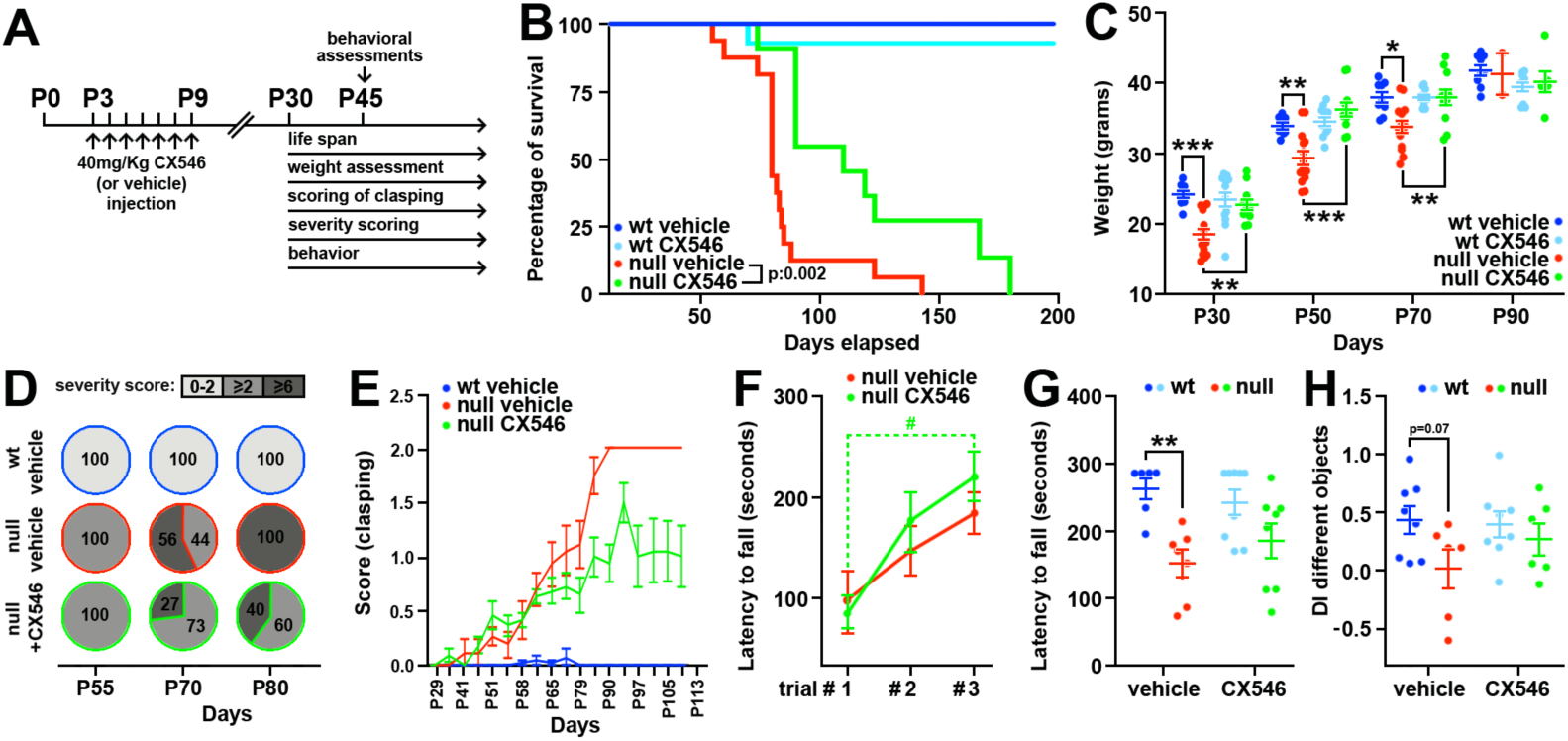
Early treatment with Ampakine ameliorates behavioral deficits of *Mecp2* null mice. **A:** Schematic representation of the *in vivo* CX546 administration protocol. **B:** Kaplan Mayer test shows an improved lifespan of *Mecp2* null CX546 treated animals (n=11) compared to *Mecp2* null control animals (n=16). **: p-value<0.01. To be noticed no difference was detected between wt mice treated with CX546 (n=14) and wt control mice (n=11). **C:** The body weight of mice was assessed at four different time points (P30, P50, P70 and P90). Two way ANOVA indicated a significant effect for both time (F(3,143)=301 and p<0,0001) and treatment (F(3,143=7,678 and p<0,0001).Tukey’s multiple comparison test was used to compare within each time point the effect of the CX546 treatment on wt and *Mecp2* null mice. At P30, P50 and P70 *Mecp2* null mice showed a statistically significant decrease in body weight compared to both wt mice treated with vehicle and wt mice treated with CX546, while *Mecp2* null mice treated with CX546 did not show any significant difference compared to the two wt groups. *: p-value<0.05; **: p-value<0.01; ***: p-value<0.005. Sample size: n=25 wt (11 treated with vehicle and 14 treated with CX546) and n= 27 ko (11 treated with vehicle and 16 treated with CX546). Each dot represents a single animal. **D,E**: The severity score, typically used in RTT phenotypic assessments (Guy *et al*, 2001; Cobolli Gigli *et al*, 2016) was used to group animals into three severity classes: absence of phenotypes (0-1,5; white), mild phenotypes (≥2-5,5; light grey), severe phenotypes (≥2-8; dark grey), as described in Patrizi *et al* (2016). The percentage refers to the number of mice falling in one of the three severity classes over the total number of animals per group. Severity scores were calculated at three different time points (P55, P70, P80). The score for hind limbs clasping was assessed twice a week starting from P30 (E). Sample size for D and E: n=25 wt (11 treated with vehicle and 14 treated with CX546) and n= 22 ko (11 treated with vehicle and 11 treated with CX546). **F:** Line graph shows mice motor learning assessed on the accelerating rotarod during three consecutive trials. Two way ANOVA indicated a significant effect of time (F(2,36)=9,9 and p=0,0004) but not of genotype. The Bonferroni multiple comparison test highlighted a significant increase of the latency to fall between trial 1 and trial 3 only for the null treated with CX546. #: p-value<0.005. **G**: The graph shows latency to fall (in seconds) on the accelerating rotarod in the last trial. Two way ANOVA indicated a significant effect of the genotype (F(1,26)=15,8 and p=0,0005). Bonferroni *post hoc* test highlighted a significant difference only between wt and ko treated with vehicle. **: p-value<0.01. Values are represented as average ± SEM. **H**: The graph shows the assessment of the discrimination index (DI) between two different objects for each mouse during the last trial of a Novel Object Recognition test. Two way ANOVA indicated a significant effect of the genotype (F(1,26)=8,763 and p=0,0065). The Bonferroni multiple comparison test highlighted a p-value of 0,07 between wt and ko treated with vehicle but not between wt and ko treated with Ampakine. Data are represented as values ± SEM. Sample size for F-G: n=15 wt(6 treated with vehicle and 9 treated with CX546) and n=15 (7 treated with vehicle and 8 treated with CX546). Sample size for H: n=17 wt (8 treated with vehicle and 9 treated with CX546) and n= 12 ko (6 treated with vehicle and 6 treated with CX546). Each dot represents a single animal.

As expected, the lifespan of CD1 KO mice treated with vehicle was markedly reduced compared to wt groups, showing 50% of survival animals at P80 (Figure 5B; Cobolli Gigli *et al*, 2016). However, when CX546 was injected in the selected early time window, mice showed a significant extension of their life-span, with 90% of animals being able to survive after P88, an age reached only by the minority of the null untreated progeny. Indeed, the average survival of CX546 treated null animals was P110 (Figure 5B).

The prolonged lifespan was accompanied by a significant improvement of the overall health of KO treated mice. First, the thoroughly described reduction in body weight of *Mecp2* null mice was markedly recovered after the treatment with CX546 at all the analyzed time points (Figure 5C). By means of the scoring system typically used to assess the worsening of the health conditions of null mice (Guy *et al*, 2001), we noticed a marked difference in the severity between untreated and treated ko animals during the overt phase of the pathology (Figure 5D). At P70, in fact, while 56% of untreated ko mice exhibited a severe RTT score (>6), only 27% of CX546 treated mice fell in this highly symptomatic range. Such difference is even more evident at P80, when we detected that all the untreated null mice fell in the highest severity score, as opposed to only 40% of the CX546 treated mice. Among the phenotypes observed during the scoring assessment, hind limb clasping, one of the classical parameter to assess neurological defects in RTT animal models, showed the beneficial effects of CX546 (Figure 5 panel E). Next, we assessed the effectiveness of the treatment in ameliorating the typical defects displayed by RTT mouse models in locomotor activity, coordination and spatial memory. Mice were tested at P45 on the accelerating rotarod test and, both the learning ability, evaluated by testing mice over the three successive trials (Figure 5F), and the typically impaired motor coordination, assessed on the last trial (Figure 5G) were significantly ameliorated by the CX546 treatment. At the same age, we tested the animals in the Novel Object Recognition test (NOR) proving that the early exposure to Ampakine improved the ability of null animals to discriminate between two different objects (Figure 5H and S5).

## Discussion

Recent preclinical and clinical studies demonstrate that besides the importance of Mecp2 in maintaining neuronal structures (Guy *et al*, 2007; Cheval *et al*, 2012; McGraw *et al*, 2011; Nguyen *et al*, 2012), its deficiency strongly affects embryonic and early postnatal development (Ip *et al*, 2018), thus long before the full outbreak of symptoms (Cosentino *et al*, 2019; Zhang *et al*, 2019). We contributed to this novel perspective by demonstrating that *Mecp2* null neurons display aberrant transcriptional profiles, reduced responsiveness to stimuli and poor morphology already during early corticogenesis (Bedogni *et al*, 2016; Cobolli Gigli *et al*, 2018). Based on the crucial role played by neuronal activity in the establishment of mature neuronal networks (Spitzer, 2006), we hypothesized that the reduced responsiveness of *Mecp2* null neurons could participate to their poor transcriptional and morphological features that would further contribute to the overall immaturity of the networks. We thus tested the rescue potential of an early enhancement of activity in *in vitro* and *in vivo Mecp2* null models.

To enhance activity, we used the Ampakine CX546, a positive modulator of AMPA glutamatergic receptors that upon glutamate binding delays deactivation and desensitization (Lynch *et al*, 2006). The choice of CX546 was based on its ability to stimulate dendritic outgrowth and maturation of *in vitro* differentiating neurons (Schitine *et al*, 2012) and was further supported by the fact that CX546 was already successfully used in preclinical studies on RTT, although in later time windows compared to this study (Ogier *et al*, 2007) and focusing on a different pathological mechanism (Degano *et al*, 2014). Our data demonstrate that the maturation of *Mecp2* null neurons is sensitive to depolarization. In fact, CX546 rescued network function, transcriptional levels and morphological complexity, thus restoring each component of the already described feed forward-cycle that ensures neuronal maturation (Spitzer *et al*, 2006). In good accordance with our hypothesis, the early treatment resulted more effective compared to the late one possibly because during early time windows neuronal networks are more plastic compared to later stages, when changes are restrained by advanced differentiation. In fact, CX546 treatment increased the levels of expression of *Gap43* and *Tuj1*, two structural proteins that are involved in synaptic plasticity and growth cones stability (Benowitz & Routtenberg 1997; Korshunova *et al*, 2010; Tischfield & Engle 2010). On the same line, CX546 exposure increased transcription of the neurotrophin *Bdnf*, a master regulator of plasticity (Lauterborn *et al*, 2003; Simmons *et al*, 2009) thoroughly associated with MeCP2 deficiency (Li *et al*, 2017; Renthal *et al*, 2018). From a functional perspective, CX546 rescued the responsiveness of null neuronal networks to stimuli, while decreasing their intracellular Cl^-^ levels, which suggests a restored responsiveness to GABA. This result fits with the recovered transcriptional levels of *Kcc2*, the neuron specific K^+^-Cl^-^ co-transporter (KCC2) that is crucial for GABA signaling maturation (Ben-Ari *et al*, 2012) and its normalization in Rett syndrome currently represents a promising therapeutic approach (Tang *et al*, 2016; Tang *et al*, 2019; Lozovaya *et al*, 2019).

Importantly, the simultaneous treatment of *Mecp2* null neurons with CX546 and Nifedipine, a voltage gated calcium channels blocker, restricted the observed transcriptional rescues, therefore proving that most of the beneficial effects were due to enhanced neuronal activity. Given the importance of calcium signaling in controlling many transcriptional programs involved in neuronal development (Rosenberg & Spitzer, 2011), our data further support the concept of a deficit of calcium-dependent maturation mechanisms in *Mecp2* null neurons (Bedogni *et al*, 2016).

A further validation to our hypothesis derived from the benefits we detected in *Mecp2* null animals treated from P3 to P9 with CX546. Indeed, we demonstrate that such a short and early treatment significantly prolonged the life span of knock out mice, ameliorating their phenotypic and behavioral scores more than 30 days after the treatment. Overall our *in vivo* results suggest a generally slower progression of the pathological symptoms together with a remarkably long-lasting effect of the treatment, thus reinforcing the role of early changes in plasticity on the functions of *Mecp2* null neuronal network. While the alteration of the already described feed-forward cycle during early development does not produce overt symptoms, such alterations likely concur to the establishment of an improper program of gene expression that takes part to the outbreak of overt symptoms and the typical *Mecp2* null network dysfunctions later in life (Nelson & Valakh, 2015; Dani *et al*, 2005; Durand *et al*, 2012; Sceniak *et al*, 2015). We thus propose the existence of an early molecular phase of Rett syndrome during which the initial developmental trajectory undertaken by maturing null networks deranges from physiology. We believe that a thorough comprehension of the mechanisms playing a pathogenic role during this phase will likely reveal novel therapeutic strategies.

The importance of targeting early neuronal maturation has already been implied by previous preclinical and clinical studies. Indeed, the injection of a truncated form of IGF1 in pre-symptomatic (P15) *Mecp2* null and heterozygous mice rescued neuronal morphology, synaptic density and network plasticity (Tropea *et al*, 2009). Further, in clinical trials, IGF1 (Trofinetide) revealed more effective when administered at pediatric ages compared to adolescence/adult (Banerjee *et al*, 2019). Similarly, the efficacy of the GABA_B_ receptor agonist Arbaclofen in the cure of Fragile-X syndrome symptoms resulted strictly dependent to the timing of treatment (Berry-Kravis *et al*, 2018). Accordingly, in a mouse model of epileptic encephalopathy, GABAergic tone modulators ensured phenotypic rescue only when administered within the first two weeks of life (Marguet *et al*, 2015).

In the next future, we will assess *in vivo* whether the time window we targeted with CX546 should be enlarged to ensure different and possibly stronger rescue effects and how such time window could be adapted for actual translational studies. Further, we will test the benefits of repeating the treatment in order to ensure the preservation, rather than the establishment alone, of functional neuronal networks.

All in all, our study suggests, as already proposed, that overt symptoms are the final outcome of a pathogenic process that likely starts very early during development, implying the importance of re-evaluating the definition of the “pre-symptomatic” phase of Rett syndrome (Cosentino *et al*, 2019; Bedogni *et al*, 2016; Zhang *et al*, 2019).

## Materials and Methods

### Animals and tissues

The *Mecp2* mouse strain was originally purchased from Jackson Laboratories (B6.129P2(C)-*Mecp2*^tm1.1Bird^/J) and transferred on a CD1 genetic background. The phenotypes affecting these animals have been previously described (Cobolli Gigli *et al*, 2016; Guy *et al*, 2001). Time pregnant females were generated by crossing overnight wt CD1 males with *Mecp2*^*-/+*^ heterozygous females; the day of vaginal plug was considered E0.5. All procedures were performed in accordance with the European Community Council Directive 86/609/EEC for care and use of experimental animals; all the protocols were approved by the Italian Ministry for Scientific Research and by the local Animal Care Committee.

### Neuroprecursors and neurons cultures

Cerebral cortices were dissected from E15.5 mouse embryos and pulled according to the genotypes. Generation and maintenance of neurosphere cultures were performed as already described (Cobolli Gigli *et al*, 2018; Figure 1A). To induce differentiation, neurospheres were dissociated in single cell suspensions and plated on matrigel-coated surfaces. Until adhesion, cells were grown in complete medium (DMEM/F12, 0.6% Glucose, 1% L-Glutamine, 1% Pen/Strep, 4µg/ml Heparin, 1x Hormone Mix, 20 ng/ml Epidermal Growth Factor (EGF), 10 ng/ml basic Fibroblast Growth Factor (bFGF)). EGF and bFGF were withdrawn respectively two and three days after plating before the addition of Fetal Bovine Serum (FBS). This last passage was considered DIV0. Generally, cells were plated at a density of 26000 cells/cm^2^. To enhance activity, CX546 (Tocris) was dissolved in DMSO and diluted 1:10 in water to obtain the working solution. The drug was added daily for 4 days directly to the medium from either DIV3 or DIV7 reaching a final concentration of 20 µM. An equal volume of 10% DMSO in water was added to controls. CX546 was washed out following the last administration by replacing the cell medium.

Primary neuronal cultures were prepared from E14.5 mouse cerebral cortices (Gandaglia *et al*, 2018). 15000 neurons/well were plated on poly-L-lysine (0,1 mg/mL; Sigma-aldrich) coated 96 multi-wells (Greiner) and cultured in Neurobasal medium supplemented with B27, L-Glutamine and Pen-Strep. CX546 (10 µM) or corresponding volume of DMSO were directly added to the medium at DIV1 and DIV3 for the early treatment and at DIV7 and DIV9 for the late treatment. Nifedipine (10 µM; Tocris) was added at the same time of CX546 only within the early time window at DIV1 and DIV3. Drugs were then washed out by changing the medium at DIV5 or DIV 11.

### Cell immunofluorescence

Cells were fixed with 4% PFA for 20 minutes, rinsed with PBS and incubated for 1 hour in blocking solution (10% horse serum, 0.1% TritonX-100). Cells were then incubated overnight at 4°C with primary antibodies diluted in blocking solution at the concentrations reported in Table S2. After PBS washing, cells were incubated for 1 hour at room temperature in secondary antibodies (1:500 in blocking solution; Molecular Probes). Before mounting, cells were rinsed with PBS and nuclei were counterstained with DAPI (Invitrogen). An upright Nikon microscope was used for acquisitions, while ArrayScan XTI HCA reader (Thermofisher) was used for all the high-throughput acquisitions.

### RNA purification, cDNA synthesis and quantitative PCR

Cells were rinsed once with PBS and lysed in Purezol (300 µL for roughly 30000 cells). RNA was isopropanol precipitated and treated with DNAse I (Sigma); next, RNA was again purified through phenol-chloroform, isopropanol precipitated over night at -20°C and RNA pellets were then dissolved in 10 µL of TE buffer. RNA quality was checked using Bioanalyzer RNA 6000 Nano chips (Agilent).

50 ng of total RNA were reverse-transcribed using Superscript IV Reverse Transcriptase (Thermo Fisher). 1 ng of cDNA for each sample was preamplified using a 0.2X pool of 74 primers (TaqMan; Thermo Fisher; Table S1) for 18 cycles using PreAmp Master Mix (Fluidigm) to enable multiplex sequence-specific amplification of targets.

Preamplified cDNAs were then diluted and assessed using a 96 X 96 qPCR Dynamic Array microfluidic chip (Fluidigm) following the manufacturer’s instructions. Baseline correction was set on Linear (Derivative) and Ct Threshold Method was set on Auto (global). To normalize, for each sample a “pseudogene” (D’haene *et al*, 2012) was obtained by averaging the Ct value of each gene depicted in Table S1 except for *Mecp2*. Each sample was then normalized against its “pseudogene”. Excel, Prism and R were used to elaborate data.

### MEA recordings

Neuroprogenitors were plated at a density of 85000 cells/well onto 12-well planar MEA chips (Axion BioSystems, Atlanta, GA), comprising 64 electrodes/well. Spontaneous extracellular activity was recorded with Axion Biosystems software (Axion Integrated Studio), by individually setting a voltage threshold for each channel equal to 7 times the standard deviation of the average root mean square (RMS) noise level. Single extracellular APs were detected by threshold crossing of 200 Hz high-pass filtered traces. Spike detection was carried out using the Axion BioSystems software NeuralMetric Tool. Evoked extracellular activity was measured after addition of 100 µM 4-AP directly to cell medium. An electrode was defined active when it registered at least 2 spikes/min. Weighted mean firing rate (WMFR) was expressed as mean number of spikes per second calculated on the active channels of the network. Bursts within single channels were identified by applying an interspike interval (ISI) threshold algorithm (Chiappalone *et al*, 2005) by defining bursts as collections of a minimum number of spikes (Nmin = 5) separated by a maximum interspike interval (ISImax = 100 ms). Only wells showing at least three active electrodes were included in the analysis.

### Automated analysis of calcium transients

Primary cortical neurons were loaded with 2 µM Fluo-4 (Invitrogen) in KRH (Krebs’–Ringer’s–HEPES containing (in mM): 125 NaCl; 5 KCl; 1,2 MgSO_4_; 1,2 KH_2_PO_4_; 25 HEPES; 6 glucose; 2 CaCl_2_; pH 7.4) for 30 minutes at 37°C and then washed once with the same solution. Stimulation was performed automatically by using the liquid handling system of the ArrayScan XTI HCA Reader (Thermo Fisher Scientific). To stimulate, one dose of NMDA (100 µM at the rate of 50 µl/sec) was added while images were digitally acquired with a high-resolution camera (Photometrics) through a 20x objective (Zeiss; Plan-NEOFLUAR 0,4NA). Hoechst fluorescence was imaged as well. 70 frames were acquired at 1 Hz with 40 msec exposure time for Fluo-4 and 25 msec exposure time for Hoechst. At least 8 baseline images were acquired before stimulation. The analysis was done with HCS Studio software using SpotDetector bioapplication (Thermo Fisher Scientific). Hoechst positive nuclei were identified and counted and the mean intensity of the Fluo-4 signal was measured in the cell body area of each cell; background intensity was measured and subtracted from the mean intensity. Only cells with neuronal morphology were included in the analysis. Calcium responses were measured as ΔF/f_0_.

### Chloride imaging recordings

Intracellular chloride measurements were performed using the N-(Ethoxycarbonylmethyl)-6-Methoxyquinolinium Bromide (MQAE) (Biotium) chloride sensor.

Briefly, cortical neurons at 14 DIV from wt and *Mecp2* null embryos were loaded with 5 μM MQAE for 45 minutes at 37°C in culture medium. Coverslips were then rinsed with external solution (Krebs’–Ringer’s–HEPES (KRH): 125 mM NaCl, 5 mM KCl, 1.2 mM MgSO_4_, 1.2 mM KH_2_PO_4_, 2 mM CaCl_2_, 6 mM glucose, and 25 mM HEPES–NaOH), pH 7.4 and transferred to the recording chamber for acquisitions.

Olympus IX81 inverted microscope, 20X dry objective (Olympus, UPLFLN NA 0.5) provided with MT20 widefield source and control system with excitation 340 nm and emission filter centered at 500 nm were used. EXcellence RT software (Olympus) was used to set up experiments and samples’ recordings. For each sample, a minimum of eight fields of interest (mean of cells observed for field, 20) have been chosen to perform live imaging recordings. Offline analyses were then made after defining ROIs centered to cell bodies and measuring MQAE mean intensity during the time recording window.

### Morphological analysis

Morphological analysis was performed on images acquired with a Nikon fluorescence microscope using a 20x objective. A binary mask was created for each Tuj1 positive cell using Adobe Photoshop by an observer blind to the genotype and the treatment. Sholl analysis was performed using the dedicated plugin of ImageJ. The smallest circle was positioned at 5 µm from the soma, while the largest at 300 µm. The distance between each circle was 10 µm. The total dendritic length (TDL) and maximal dendritic length (MDL) were evaluated using NeuronJ, a plugin of ImageJ. For each neuron, the length of each dendrite (Lm) was measured and the sum of such values defined the total dendritic length (TDL). The polarization value was calculated as the ratio between the maximal dendritic length and the average dendritic length. A ratio equal or above 2 defines a polarized neuron; a culture is defined polarized when at least 60% of neurons within it are polarized (Horton *et al*, 2006).

### Cell viability assay

Cell viability was assessed through MTT assay (3-(4,5-dimethylthiazol-2-yl)-2,5-diphenyltetrazolium bromide; Sigma). MTT (4 mg/mL) was diluted 1:10 in DMEM-F12 and added to each well after medium withdrawal; plates were then incubated at 37°C for 4 hours. Next, the solution was replaced by 100 µL of DMSO and the colorimetric signal assessed with a spectrophotometer (570 nm).

### *In vivo* Drug administration

Ampakine CX546 40 mg/kg was administered daily via a subcutaneous injection for seven days (starting from P3) at the same time. The drug was solubilized in ethanol (100%) and diluted 1:100 in saline to obtain the working concentration. For the vehicle treatment, ethanol (100%) was diluted 1:100 in saline. A total of eight litters were treated either with Ampakine or vehicle, genotype and treatment were blind to the operator.

### Neurobehavioral Characterization and phenoptypic scoring

Weight and phenotypic scoring assessment were evaluated twice a week starting from P30. The phenotypic assessment was performed as described previously (Cobolli gigli *et al*, 2016). Scores were classified in three different groups of severity as in Patrizi *et al*, 2016. To be noticed, mice that rapidly lost weight were euthanized for ethical reasons. To avoid any bias, an investigator blind to the genotypes of the tested animals performed all the analyses.

### Behavioral assessment

Animals were maintained on an inverted 12h light/darkness cycle at 22-24°C. A total of 18 wt animals and 14 ko animals were involved in the behavioral assessment.

#### Rotarod

Mice were assessed on an accelerating rotarod (Ugo-Basile, Stoelting Co.). The test was carried out in three days, two days of training and one day of test. Each session consists of three trials and each trial lasts five minutes. Revolutions per minute (rpm) were set at an initial value of 4 with a progressive increase to a maximum of 40 rpm. Each trial ended when the mouse fell down or after 5 minutes.

Latency to fall was measured by the rotarod timer. Data were plotted and subjected to statistical analysis with Graphpad Prism.

#### Novel Object Recognition (NOR)

The test was performed in an open square arena of 50×50 cm and consists of three different sessions. On day 1, during the first session, mice were allowed to habituate with the arena for 10 minutes. On day 2, mice underwent the training phase (5 min) in which they were allowed to explore two identical objects. Object investigation was defined as time spent sniffing the object when the nose was oriented toward the object and the nose-object distance was 2 cm or less. Time spent sniffing two identical objects during the familiarization phase confirmed the lack of an innate side bias. The total time spent by the mice to explore the two objects in the cage was also measured; if this was less then 30s (10% of the total time) the mouse was not included into the final test (Figure S5). Finally, on day 3, mice were tested (5 min) using two different objects. Recognition memory, defined as spending significantly more time sniffing the novel object than the familiar object, was calculated as Discrimination Index (DI): difference between the exploration time for the novel object and the familiar object, divided by total exploration time. The sessions were recorded with the video tracking software Ethovision XT (Noldus).

### Statistical analysis

Statistical analysis and plotting of data were performed with Graphpad Prism. Student’s t test was used for the statistical analysis when wt were compared to Mecp2 null mutant samples. One-way ANOVA was used to statistically compare the effects of early and late addition of CX546 or the effect of CX546 or CX546 plus Nifedine on *Mecp2* null neurons. Data from the experiment on wt and null NPCs or neuronal cultures at different DIVs, from the assessment of mice weight and from mice behavioral experiments were analyzed by two-way ANOVA. When there was a significant effect of treatment or genotype, or a significant interaction between the variables (one way or two-way ANOVA), appropriate post-hoc test was applied. A p value of 0,05 was considered significant. Possible outliers within an experimental group were identified with Grubb’s test.

## Figure legends

**Figure S1:**
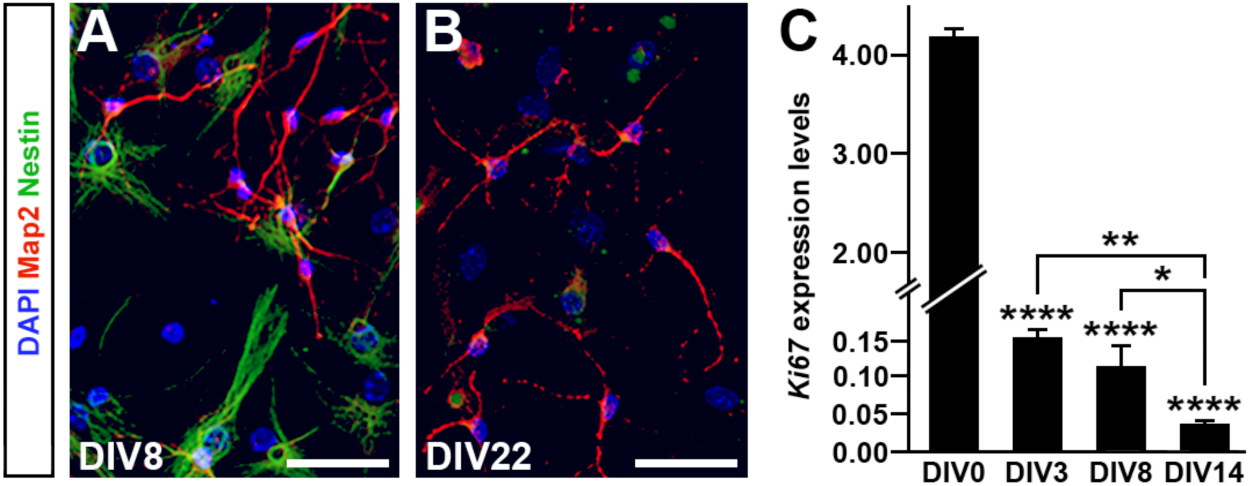
Cell proliferation is absent in DIV 22 differentiated cultures. To investigate the presence of proliferating cells at the end of the differentiation protocol cultures were stained with Nestin (marker of neuroprogenitors) and Map2 (marker of neurons) at DIV8 and DIV22 (panels A and B); *Ki67* (marker of cell cycle) expression levels (2^-Δct^) at different DIVs were also measured. The absence of Nestin positive cells at DIV22 and the decreased expression of *Ki67* through time indicated the progressive completion of the differentiation scheme of the cell types composing the cultures. Panel C: Two-Way ANOVA and Bonferroni post-hoc test. **: p-value<0.01; ****: p-value<0.001. Histograms show the mean ± SEM from at least 6 wells obtained from two independent preparations. Scale bar A,B: 20 µm.

**Figure S2:**
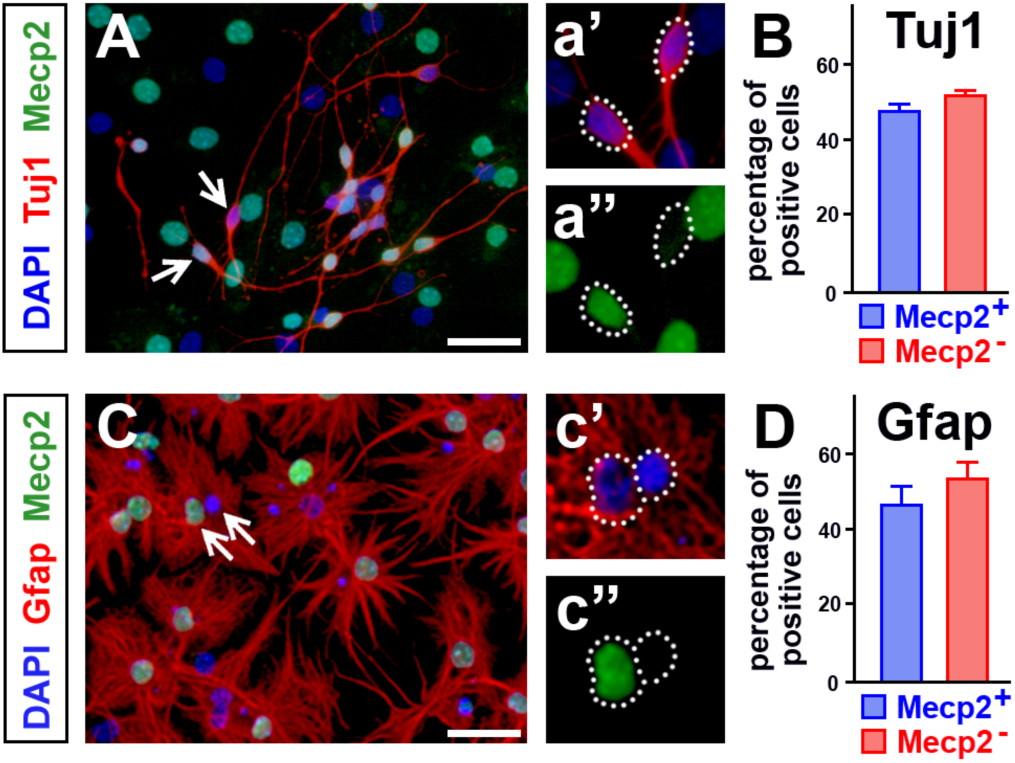
Lack of Mecp2 does not influence the fate commitment of NPC deriving from heterozygous cortices. Nuclei deriving from heterozygote cortices were stained for Mecp2 and DAPI, neurons and astrocytes were distinguished using antibodies against Tuj1 and GFAP respectively. The percentage of cells positive or negative for Mecp2 was counted at DIV22 over the total of Tuj1 (B) and GFAP (D) positive cells. No difference between the two groups was highlighted (Student’s t-test). Histograms show the mean ± SEM from at least 3 samples obtained from two independent preparations. Scale bar: 30 µm.

**Figure S3:**
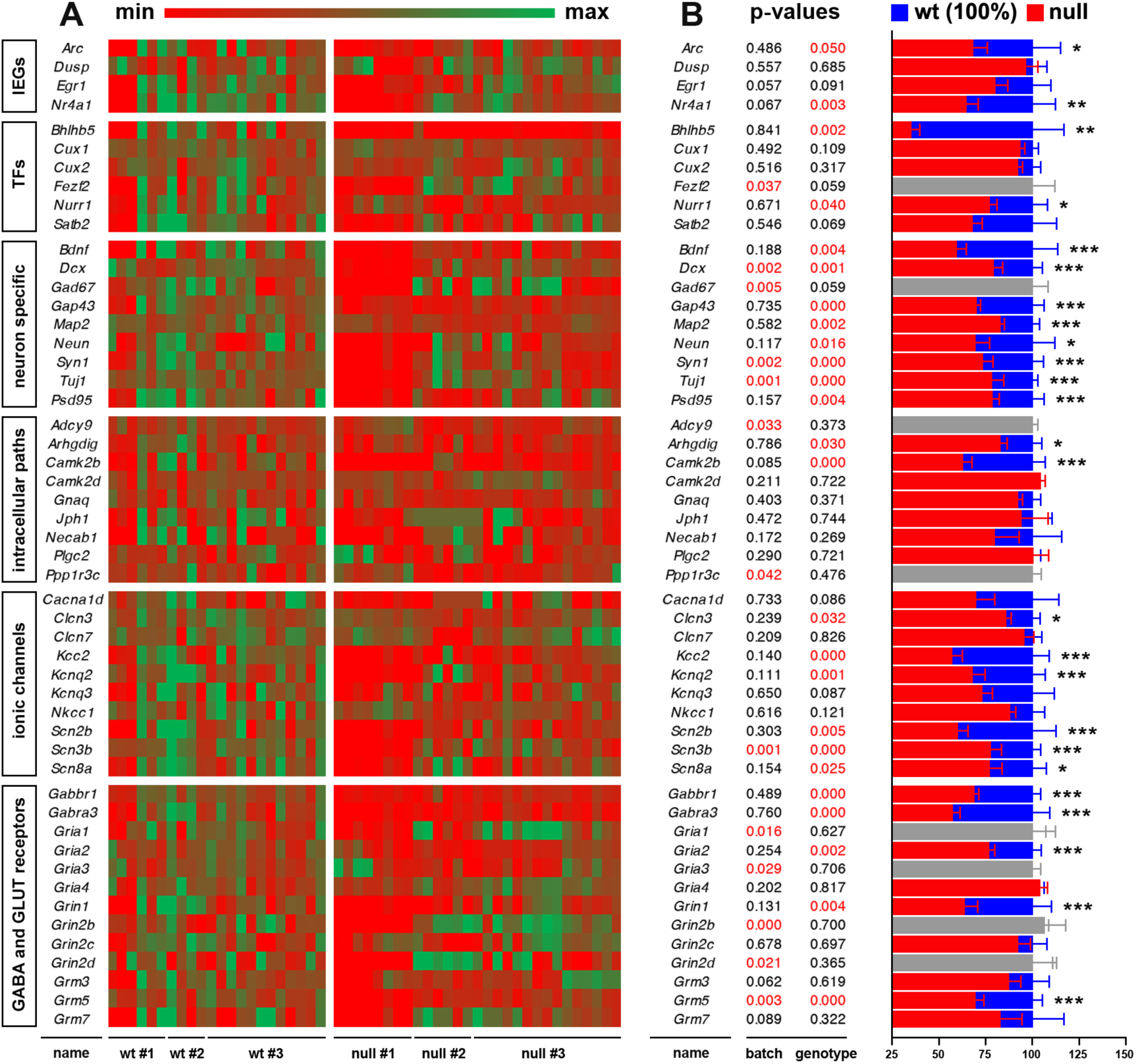
Transcriptional differences between wt and *Mecp2* null samples in basal condition. **A**: Heat map of the levels of 51 transcripts expressed by post-mitotic neurons in three distinct batches (#1, 2 and 3) per genotype. Genes were divided in clusters depending on their biological function. IEGs: Immediate early genes; TFs: transcription factors. Each batch was composed at least by four samples. **B**: Two-way ANOVA followed by Bonferroni *post hoc* test, in which the variables were genotype and batch, was used to highlight and discard batch effects. Our statistical approach identified 43 transcripts that survived the statistical cutoff, while only 8 were discarded (depicted in grey in the graph in B). Significant p-values (<0,05) are depicted in red. Each bar of the graph represents the average expression levels for each gene composing the six clusters. Histograms show the mean ± SEM.*: p-value<0.05; **: p-value<0.01; ***: p-value<0.005.

**Figure S4:**
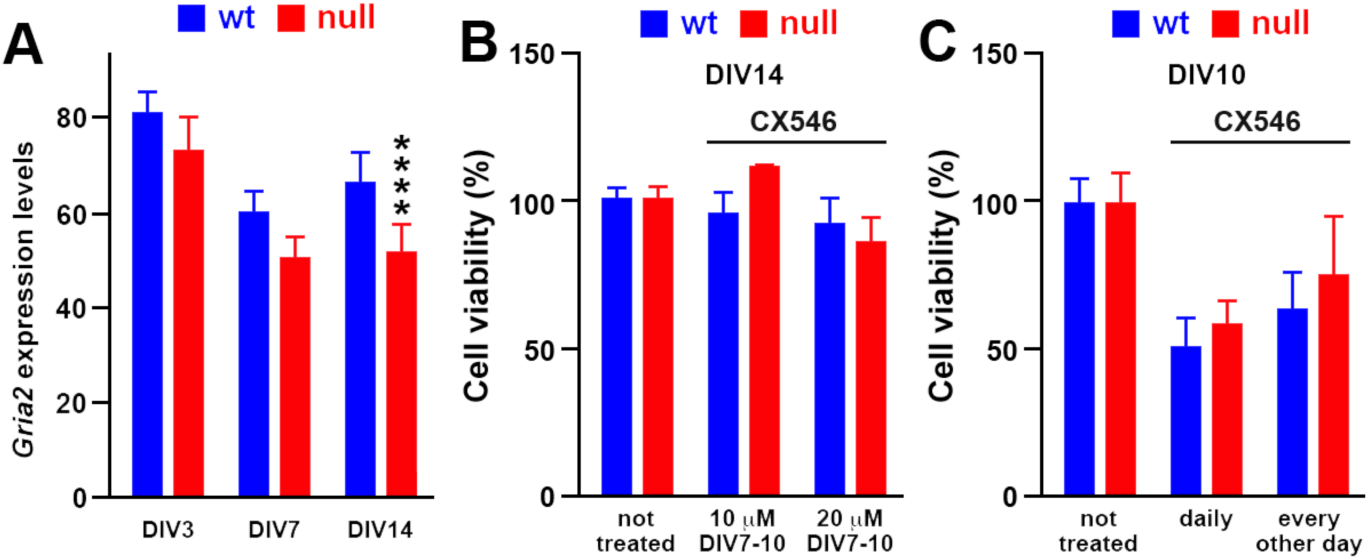
Experimental setting of *in vitro* Ampakine treatment. **A:** Bar graphs showing the expression of the AMPA receptor subunit *Gria2* mRNA in cultures differentiating from NPCs at three different time points (expressed as 2^-Δct^). Two-way ANOVA followed by Bonferroni *post hoc* test indicated a reduced expression of the gene in null neurons only at DIV14. *: p-value<0.05; ***: p-value<0.005. **B,C** To assess Ampakine CX546 toxicity we measured cell viability through MTT assay. In B cells were collected at DIV 14 after Ampakine exposure (10 and 20 µM) from DIV7 to DIV10. In C cells were collected at DIV 10 after the daily or the other daily exposure to 20 µM CX546. Two way ANOVA did not indicate any significant effect of the late (DIV7-10) CX546 treatment (B) while both the exposures in C showed a significant effect (F(2,17)=8,913 and p=0,0023), thus indicating the toxicity of this protocol. Sample size: n≥3 wells deriving from two independent preparations.

**Figure S5:**
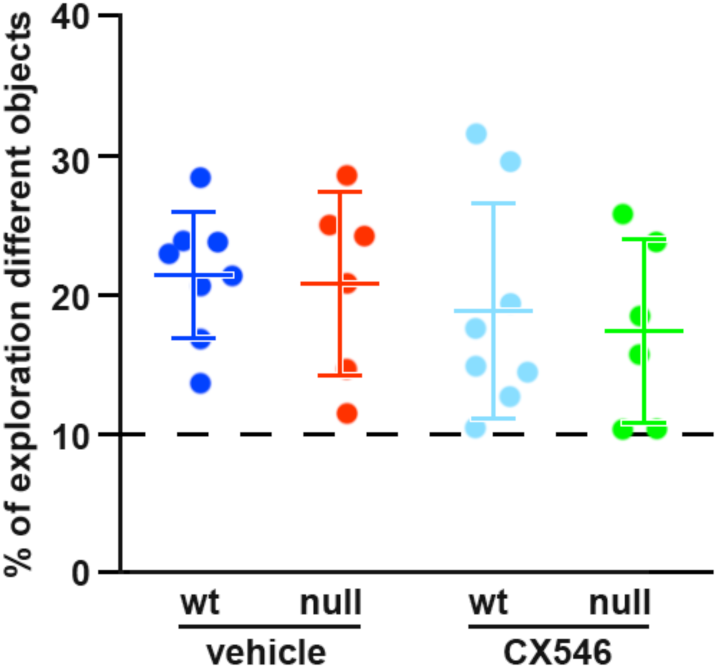
Exploration rate assessment during NOR test. The graph shows the time mice spent exploring the two objects during the third day of test (depicted as %). Two-Way ANOVA did not indicate any significant effect of both genotype and treatment. Since all mice showed an exploration rate higher then 10% of the total time, they were all included in the Discrimination index assessment (see materials and methods). Data are represented as values ± SEM. Sample size for: n=16 wt (8 treated with vehicle and 8 treated with CX546) and n=12 ko (6 treated with vehicle and 6 treated with CX546). Each dot represents a single animal.

**Table S1:**
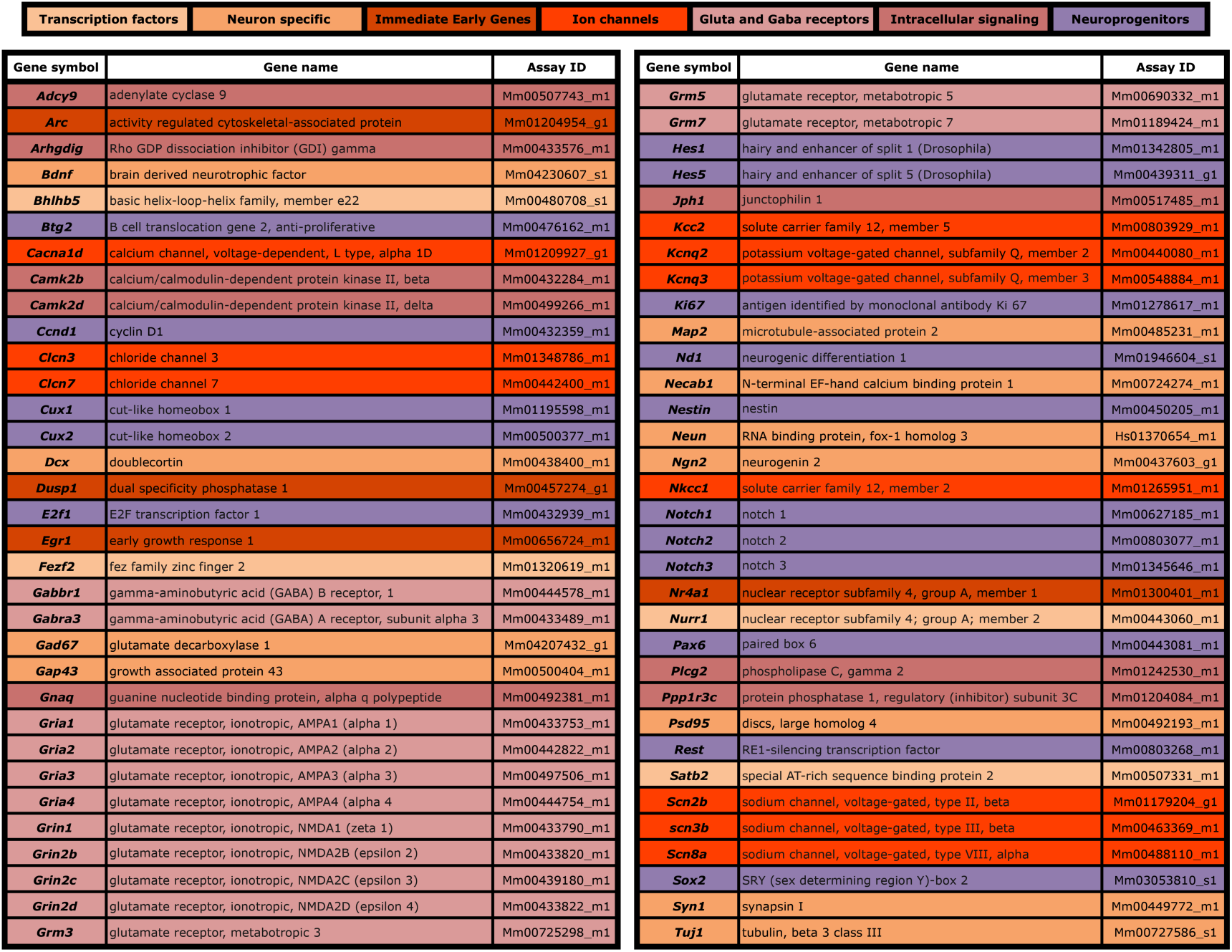
TaqMan probes classified based on biological functions; the Assay ID for Mecp2 was Mm01193537_g1.

**Table S2:**
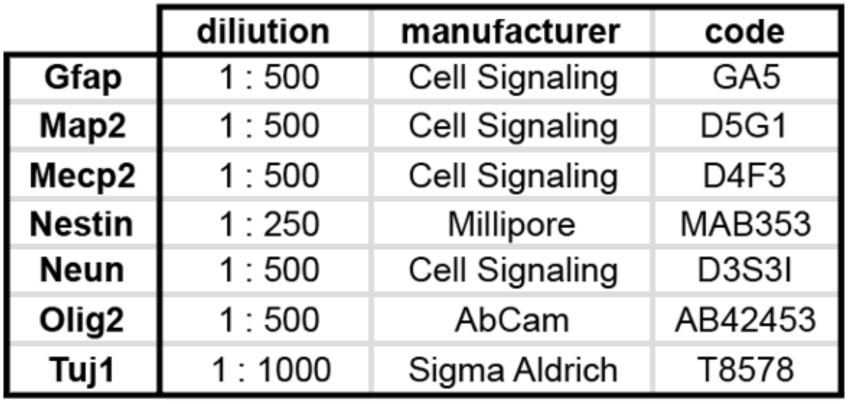
List of antibodies and dilutions.

## Acknowledgment

We are grateful to ProRett Research, an Italian association that daily provides us with both strive and funding to carry on our study on Mecp2 and Rett syndrome. Lejeune Foundation (Paris, France) and Fondazione Umberto Veronesi (Milan, Italy) provided additional funding to FBedo. AIRC (Associazione Italiana per la Ricerca sul Cancro; Grant n° IG2016-ID18575) and ERC (European Research Council, Consolidator Grant n° 617978) provided additional funding to MP. Compagnia di San Paolo Torino (9344), Ministero della Salute Ricerca Finalizzata (GR-2016-02363972) and EU Era-Net Neuron 2017 “Snareopathies” provided additional funding to FBen. The team of the microscopy facility at the San Raffaele Hospital (ALEMBIC) produced all the imaging data described in this study.

## Conflict of Interest

None declared.

